# The aerial epidermis is a major site of quinolizidine alkaloid biosynthesis in narrow-leafed lupin

**DOI:** 10.1101/2023.03.20.532575

**Authors:** Karen Michiko Frick, Marcus Daniel Brandbjerg Bohn Lorensen, Eddi Esteban, Asher Pasha, Alexander Schulz, Nicholas James Provart, Christian Janfelt, Hussam Hassan Nour-Eldin, Fernando Geu-Flores

## Abstract

Lupins are promising legume crops that accumulate toxic alkaloids in the seeds, complicating their use as high-protein crops. The alkaloids are synthesized in green organs (leaves, stems, and pods) and a subset of them is transported to the seeds during fruit development. The exact sites of biosynthesis and accumulation remain unknown, however mesophyll cells have been proposed as sources, and epidermal cells have been suggested as sinks. We examined the spatial localization of the alkaloids in biosynthetic organs of narrow-leafed lupin using mass spectrometry-based imaging (MSI). The alkaloids that accumulate in seeds (“core” alkaloids) were evenly distributed across tissues, however their esterified versions accumulated primarily in the epidermis. In addition, we generated a tissue-specific RNAseq dataset of biosynthetic organs using laser-capture microdissection. The dataset revealed that alkaloid biosynthetic genes are strongly expressed in the epidermis. To confirm the biosynthetic capacity of the leaf epidermis, we combined precursor feeding studies with mass spectrometry imaging, which showed that the lower epidermis is highly biosynthetic. Our work challenges the current assumptions on the precise sites of lupin alkaloid biosynthesis, with direct implications for the elucidation of the alkaloid biosynthesis pathway and the long-distance transport network from source to seed.

## 1. Introduction

Plants produce an immense array of specialized metabolites with important applications in agricultural, pharmaceutical, and food industries. The biosynthesis and accumulation of plant specialized metabolites is often under strict control, being synthesized in specific cell-types/tissues and often stored in others. For example, in Solanaceae, the first steps of tropane alkaloid biosynthesis occurs in the root pericycle, intermediate steps occur in the root endodermis and outer cortex, and final steps once again occur in the root pericycle (Nakajima and Hashimoto, 1999; Suzuki et al., 1999; Suzuki et al., 1999). The tropane alkaloids are then transported from the roots to the aerial parts where they are stored. In turn, the biosynthesis of monoterpene indole alkaloids (MIAs) in leaves/stems of *Catharanthus roseus* involves four types of cells: early steps occur in the internal phloem-associated parenchyma, intermediate steps take place in epidermal cells, and final steps occur in idioblast or lacticifer cells, where most of the MIAs also localize (St-Pierre et al., 1999; Yamamoto et al., 2016). A thorough understanding of the biosynthesis and accumulation of plant specialized metabolites is crucial for understanding their function, and for altering their production and/or accumulation in native or heterologous hosts.

Quinolizidine alkaloids (QAs) are a class of toxic specialized metabolites found in lupins (*Lupinus* spp.) and other Genistoid legumes. Four lupin species are currently cultivated for their high-protein seeds: narrow-leafed lupin (NLL; *L. angustifolius*), white lupin (*L. albus*), yellow lupin (*L. luteus*), and Andean lupin (*L. mutabilis*). Of these, NLL is the most widely cultivated (Vishnyakova et al., 2021). QAs can accumulate to high levels in the seeds (up to ~3%), and even the low-QA varieties (“sweet” varieties) often exceed the industry threshold to be used as food and feed (0.02% in Australia and New Zealand; (Schedule 19, 2015)). Interestingly, it has been shown that the QAs in the seeds are not made *in situ*, but are transported there from maternal tissues (Otterbach et al., 2019), likely via the phloem (Lee et al., 2006). However, it is not well understood where QAs are made in the maternal tissues, and neither is the long-distance QA transport pathway from source to seed.

Studies performed in the 1980s provided foundational knowledge on where QAs are made and accumulated in lupin plants. The first enzyme of the QA biosynthesis pathway, lysine decarboxylase (LDC) was found to localize to the chloroplast stroma in *L. polyphyllus* leaves (Wink and Hartmann, 1982). Examination of the distribution of QAs in *L. polyphyllus* petiole sections using laser desorption mass spectrometry (LAMMA 1000) showed that the QA lupanine was only detected in the epidermis and the neighboring 1-2 subepidermal layers (Wink et al., 1984). To find out in which tissues QAs are made, a radiolabeled QA intermediate (cadaverine) was fed to isolated petiole epidermis, to the corresponding “mesophyll” (petiole without the epidermis), and to intact petioles of *L. polyphyllus*. Incorporation of the intermediate into lupanine was detected in the “mesophyll” and intact petioles, but not in the isolated epidermis (Wink and Mende, 1987). The authors noted that epidermal cells are usually devoid of chloroplasts, and thus, they hypothesized that QAs are made in chloroplast-containing mesophyll cells. It was also hypothesized that QAs are transported from mesophyll to epidermal cells, which is consistent with the observed epidermal localization of lupanine as well as the high capacity of the isolated epidermis to take up externally supplied lupanine (Wink and Mende, 1987).

Decades later, the gene encoding LDC was identified in NLL (Bunsupa et al., 2012), and so was the gene encoding the second enzyme in the pathway, copper amine oxidase (CAO) (Yang et al., 2017). Examination of *LDC* and *CAO* expression in whole NLL organs revealed that they are highly expressed in green parts of the plant—leaves, stems and developing pods (Bunsupa et al., 2012; Yang et al., 2017; Frick et al., 2018). As expected, fluorescently tagged LDC was found to localize to plastids (Bunsupa et al., 2012). These studies seem to support the hypothesis that QAs are made in the “mesophyll”, or rather, in the chloroplast-containing parenchyma of the above-mentioned organs. It follows that QAs are transported from there to the epidermis.

Here, we set out to determine the exact sites of QA accumulation and biosynthesis in the QA-producing organs of NLL (leaf, stem, and developing pod). In our studies, we used mass spectrometry-based imaging (MSI), laser-capture microdissection coupled to RNAseq, and precursor feeding studies coupled to LC-MS and MSI. Our results indicate that the epidermis (and not the mesophyll) is the major site of biosynthesis, leading to a new spatial model for QA biosynthesis and transport.

## 2. Materials and Methods

### 2.1. Plant growth conditions

Seeds of NLL cv. Oskar (a high-QA cultivar obtained from Hodowla Roślin Smolice, Poland) were sown in 16 cm-wide, 20 cm-deep pots at a density of two seeds per pot in potting mix (Pindstrup Færdigblandig 2; www.pindstrup.dk). The plants were grown in a growth chamber at day/night temperatures of 20/18 °C, a light/dark photoperiod of 16/8 h, and 60 % relative humidity. Light was supplied at 220 μmol/m^2^/s during the day.

### 2.2 Preparation of tissue for cryosectioning

Tissue (leaf, stem, and whole pods with seeds) was collected from plants when the first pods reached 30 days after anthesis (DAA). The tissue was quickly cut into 1.5–3 cm length pieces, frozen in liquid nitrogen, and stored at −80 °C until further analysis. Frozen tissue was freeze-embedded as described before (Kawamoto and Kawamoto, 2021), but without tissue fixation. Briefly, embedding containers obtained from SECTION-LAB Co. Ltd., Japan (www.section-lab.jp) were filled with 2.5 % (w/v) aqueous carboxymethylcellulose (CMC) solution. The filled containers were pre-chilled by partially submerging in a mixture of hexane and dry ice. Frozen tissue was placed and orientated in the CMC and the whole block was fully submerged in the hexane/dry ice mixture to completely freeze it without the tissue thawing. Embedded blocks were stored at −80 °C. Leaf and stem tissues were embedded in a 2 x 1.5 cm container, while pod tissue was embedded in a 3.5 x 2.5 cm container.

### 2.3 Metabolite imaging by MALDI-MSI

CMC embedded leaf, stem, and pod tissues were transversely sectioned in a Leica CM3050S Cryostat at −30 °C using Kawamoto’s film method (Kawamoto and Kawamoto 2021) adapted for plant samples (Montini et al., 2020) to obtain 12–14 μm sections. Briefly, adhesive cryofilm (type 3C [16UF] 2 cm; SECTION-LAB Co. Ltd., Japan) was used to obtain intact sections which were then mounted onto glass slides using double-sided carbon tape (SPI supplies; 2spi.com). The sections were freeze-dried under vacuum overnight. 2,5-dihydroxybenzoic acid (DHB) matrix was sublimated on leaf and stem sections, while for pod sections it was applied by spraying. Sublimation was performed on a custom-built sublimator at a temperature of 140 °C, and spraying deposition was performed by spraying 300 μL of a 30 mg/mL DHB solution in methanol/water (90:10) using a custom-built matrix sprayer, as described in detail elsewhere (Wenande et al., 2017).

After deposition of the matrix, the samples were imaged on a Thermo QExactive Orbitrap mass spectrometer, equipped with an AP-SMALDI5 ion source (TransMIT GmbH, Giessen, Germany). Imaging was performed in positive ion mode using a scan range of *m/z* 150–1050. QAs were identified based on the accurate *m/z* ratios of their protonated forms in accordance to previous assignment in NLL plant organs using LC-MS (Otterbach et al., 2019; Mancinotti et al., 2021). Phosphatidylcholine (34:2)—a component of cell membranes–was chosen as a control metabolite for imaging, which could be visualized across the whole area of the tissue sections (data not shown). The complete list of *m/z* values for the QAs, L-lysine, their isotopically labelled counterparts, and phosphatidylcholine (34:2) is available as supplementary information (Table S1). Each sample type was analyzed by MALDI-MSI in duplicates, with similar results between the replicates (replicate images not shown). Images were generated using the software, MSiReader 1.02 (Robichaud et al., 2013; Bokhart et al., 2017). Images were normalized by the total ion current (TIC) and the color scales were adjusted to best display the compounds of interest.

### 2.4 Tissue-specific transcriptome analysis by LCM-RNAseq

CMC-embedded leaf, stem, and pod tissues were sectioned under the same conditions as for MALDI-MSI (see above), but without the use of cryofilm, and this time obtaining 14–20 μm sections. Whilst still at −30 °C, the CMC embedding media was carefully removed from the sectioned tissue as much as possible. The sections were thaw-mounted onto SuperFrost Plus Slides (VWR; treated with RNAseZAP and dried) by very briefly warming the slide with the hand to thaw the section and immediately re-freezing it on the cold inductive plate inside the cryostat (tissue was thawed for maximum 1–2 s). Mounted sections were freeze dried under vacuum overnight.

Laser-capture microdissection (LCM) was carried out on a PALM-Microbeam instrument (Zeiss) at room temperature. AutoLPC setting was used to manually collect cells (energy=100, focus=80). Whenever target cells were in close proximity to other cell types, the cut setting (energy=100, focus=80) was used prior to the AutoLPC capture of cells to remove the undesired cell types. Cells were collected in a 0.2 ml PCR tube cap filled with 25 μl RNAlater (Thermo Fisher Scientific), for a maximum time of 45 min. Immediately after collection, the first step of RNA extraction was carried out by adding 100 μl of extraction buffer from the Arcturus PicoPure RNA Isolation Kit (Thermo Fisher Scientific) and incubating samples for 30 min at 42 °C. Cell extracts were stored at −80 °C before completing the rest of the PicoPure RNA isolation protocol, adjusting for the modified volume of cell extract. The quality and quantity of RNA was determined using an Agilent RNA 6000 Pico Kit (Agilent Technologies). For each cell sample type, RNA from several extractions was pooled together until 10 ng was obtained for one sample. Due to the laborious nature of collecting cells using LCM, two biological replicates of each sample type were collected (using tissue collected from separate plants). For all RNA samples collected from pods, RIN values were >7. For RNA samples collected from leaves, RIN values were >6.5. For RNA samples collected from stems, RIN values were >7, except for parenchyma samples (5.5 and 5.7) and one phloem sample (6.5). RNA samples were provided to Macrogen (South Korea) for library construction and sequencing. Libraries were prepared using SMART-Seq™ Ultra™ Low Input RNA kit and sequenced on an Illumina platform (HiSeq X) to generate 151-bp paired-end reads.

Raw Illumina reads were pre-processed using rCorrector (Song and Florea, 2015), and TRIMGalore! (v0.6.4) to trim adapter sequences and low quality bases (<5). Any reads mapping to the SILVA small and large subunit ribosomal RNA database (for Archaeplastida; release 138.1), were removed, as well as overrepresented sequences as determined by FastQC (v0.11.9). Transcript expression of processed RNAseq data was quantified using Salmon (Patro et al., 2017) with the coding sequences from the NLL cv. Oskar transcriptome (Yang et al., 2017) as an index. Transcripts per million (TPM) values were cross-sample normalized between all samples (TMM normalization). Differential gene expression analysis was performed using DESeq2 (Love et al., 2014).

### 2.5 Generation of a NLL eFP Browser

The eFP Browser framework described in Winter et al., (2007) was modified to accept the NLL gene identifiers from the existing NLL cv. Oskar transcriptome (Yang et al., 2017). Previously obtained RNAseq data (Yang et al., 2017) and our new LCM-RNAseq data were databased on the Bio-Analytic Resource for Plant Biology (BAR) server at bar.utoronto.ca (Toufighi et al., 2005). Images representing the samples from which RNA was isolated (whole plant organs as well as microdissected leaf, stem and pod tissues) were generated using GIMP (www.gimp.org). XML files linking the appropriate regions of the images to their respective database samples were manually created in a text editor and were loaded into the Lupin eFP Browser (available at https://bar.utoronto.ca/efp_lupin/cgi-bin/efpWeb.cgi).

### 2.6 Analysis of QA biosynthetic gene expression by qPCR

Leaf abaxial epidermis was peeled from fresh NLL leaves, similarly to the method described by Weyers and Travis (1981). Peeled tissue was frozen immediately in liquid nitrogen and stored at −80 °C. The remaining leaf tissue (without the abaxial epidermis) was also collected. RNA was extracted from ~30 mg of abaxial epidermis or the remaining leaf tissue using the Spectrum Plant Total RNA kit (Sigma-Aldrich) including on-column DNase I digestion (Sigma-Aldrich). cDNA synthesis and qPCR to determine the expression of *LDC* and *CAO* was carried out as described previously (Mancinotti et al., 2021).

### 2.7 Confocal microscopy imaging of chloroplasts

At ~30 DAA, leaf, stem, and pod tissues were dissected into thin sections including the epidermis, and mounted with perfluorodecalin for observation by a SP5-X confocal laser scanning microscope equipped with a DM6000 microscope (Leica, Germany). Chlorophyll autofluorescence was visualized using excitation/emission wavelengths of 496/661–750. Images were subsequently acquired and processed using the microscope imaging software Leica Application Suite X (version 3.7.5.24914). Each sample type was imaged with three biological replicates, with similar results between the replicates (replicate images not shown). Z-stack images were also collected to visualize chloroplasts in the epidermis and underlying cell layers.

### 2.8 Feeding of detached leaves with isotopically labelled L-lysine

Leaves of 4–6 week old NLL plants were detached by cutting at the petiole, and detached leaves were fed continuously through the petiole with an aqueous solution of either 5 mM isotopically labelled L-lysine (L-lysine·2HCl [^13^C_6_, ^15^N_2_], Cambridge Isotope Laboratories, Inc.) or 5 mM unlabeled L-lysine (Sigma-Aldrich). Leaves were fed in a growth chamber at the above-mentioned conditions for 0, 6, 10, 18, and 24 h. Due to the day-night cycle in the growth chamber and the timing of the experiment, leaves fed for up to and including 10 h were subjected to uninterrupted light, whereas leaves fed for 18 and 24 h experienced a night cycle. Leaf tissues were harvested at each ot the mentioned time points. First, the abaxial epidermis was peeled and collected. Some of the remaining leaf tissue without abaxial epidermis was also collected, and some was used to harvest the mesophyll. For the latter, remaining leaf tissue was attached to plastic slides with the adaxial epidermis facing down using a minimal amount of CMC embedding media. After freezing on dry ice, a scalpel blade was used to scrape off and collect mesophyll cells, carefully avoiding the adaxial epidermis. After harvesting, all tissues were immediately submerged in liquid nitrogen and stored at −80 °C until further analysis.

### 2.9 Analysis of fed leaves by LC-MS

Tissue was pulverized with a steel ball homogenizer and metabolites were extracted from the powder using 100 μl extraction solvent (60 % methanol in water, 0.06 % formic acid, and 5 ppm caffeine as internal standard) per 3 mg of tissue. The mixtures were shaken vigorously for 90 min at room temperature after which they were centrifuged at maximum speed for 1 minute. The supernatant was collected and diluted 1:5 with water. Each diluted extract was passed through a 0.22 μm filter and transferred to a glass vial for LC-MS analysis.

LC-MS analysis was performed on a Dionex UltiMate 3000 Quaternary Rapid Separation UHPLC+ focused system (ThermoFisher Scientific). Separation was achieved on a Kinetex^®^ 1.7-μm C18 column (100 × 2.1 mm, 1.7 μm, 100 Å; Phenomenex, Torrance, CA, USA), including a SecurityGuard™ C18 guard column (2.1 mm, 2 μm; Phenomenex). Mobile phases A and B consisted of, respectively, 0.05% formic acid in water and 0.05% formic acid in acetonitrile. Analytes were eluted using the following gradient at a constant flow rate of 0.3 mL/min: 0–0.5 min, 2% B (constant); 0.5–2.375 min, 2–6% B (linear); 2.375–7 min, 6–25% B (linear), 7–13 min, 25–100% B (linear); 13–14 min, 100% B (constant); 14–14.5 min, 100–2% B (linear); and 14.5–20 min, 2% B (constant). The injection volume of all samples was 2 μl. The UHPLC was coupled to a Compact micrOTOF-Q mass spectrometer (Bruker, Bremen, Germany) equipped with an electrospray ion source (ESI) operated in positive mode. Mass spectrometer conditions were as described before (Otterbach et al., 2019; Mancinotti et al., 2021).

The complete list of *m/z* values for the target compounds and their isotopically labelled counterparts is available as supplementary information (Table S1). Labelled and unlabeled QAs were quantified based on the relative abundance of their protonated molecular ions (peak areas from extracted ion chromatograms ± 0.005 Da) compared to the internal standard caffeine. L-lysine was found to deaminate readily under analysis conditions; therefore, the relative abundance of both labelled and unlabeled L-lysine was calculated based on the deaminated fragment (Table S1). At each time point, tissues from three fed leaves were analyzed (biological replicates). However, one of the leaves collected after 10 h did not seem to be actively synthesizing QAs, and thus, only two replicates were analyzed for this time point. In addition, two biological replicates were analyzed for abaxial epidermis tissue collected at 24 h after feeding due to the loss of one of the samples. For statistical comparisons, data that did not follow a normal distribution was log-transformed. For data following a normal distribution (prior or after log-transformation), one-way ANOVA was used to assess statistical differences between the relative abundance of detected compounds between tissues (independently for each time point). A significant one-way ANOVA was followed up by a Tukey test for multiple comparisons (95% confidence interval). For data that did not follow a normal distribution after log-transformation, a Kruskal-Wallis test was used to determine statistical differences. A significant Kruskal-Wallis test was followed up by a Dunn’s test for multiple comparisons. A single replicate for tissue collected from leaves fed with unlabeled L-lysine was also analyzed to ensure there was no background signal for the *m/z* values of the isotopically labelled QAs (data not shown).

## 3. Results

### 3.1 Localization of QAs in biosynthetic organs by mass spectrometry-based imaging

To determine the localization of QAs in biosynthetic organs of NLL, we prepared cross sections of leaves, stems, and developing pods, and analyzed them via high-resolution matrix-assisted laser desorption/ionization-MSI (MALDI-MSI). We examined the distribution of eight QAs, including core QAs representing unmodified or minimally modified QA backbones (lupanine, 13-hydroxylupanine, and angustifoline) as well as esterified QAs representing larger modifications resulting from conjugation to CoA-activated acids (e.g. 13-*cis*/*trans*-cinnamoyloxylupanine).

#### 3.1.1 Leaf

As the NLL leaf is bilaterally symmetrical, we imaged one half of the leaf including the central midvein (Fig. 1A). The core QAs lupanine and angustifoline appeared generally evenly distributed throughout the leaf, while 13-hydroxylupanine appeared to have a higher distribution towards the midvein (Fig. 1, B-D). Interestingly, the esterified QAs showed a different distribution pattern and appeared to be most highly localized in the epidermis (Fig. 1, E-H).

**Fig. 1:**
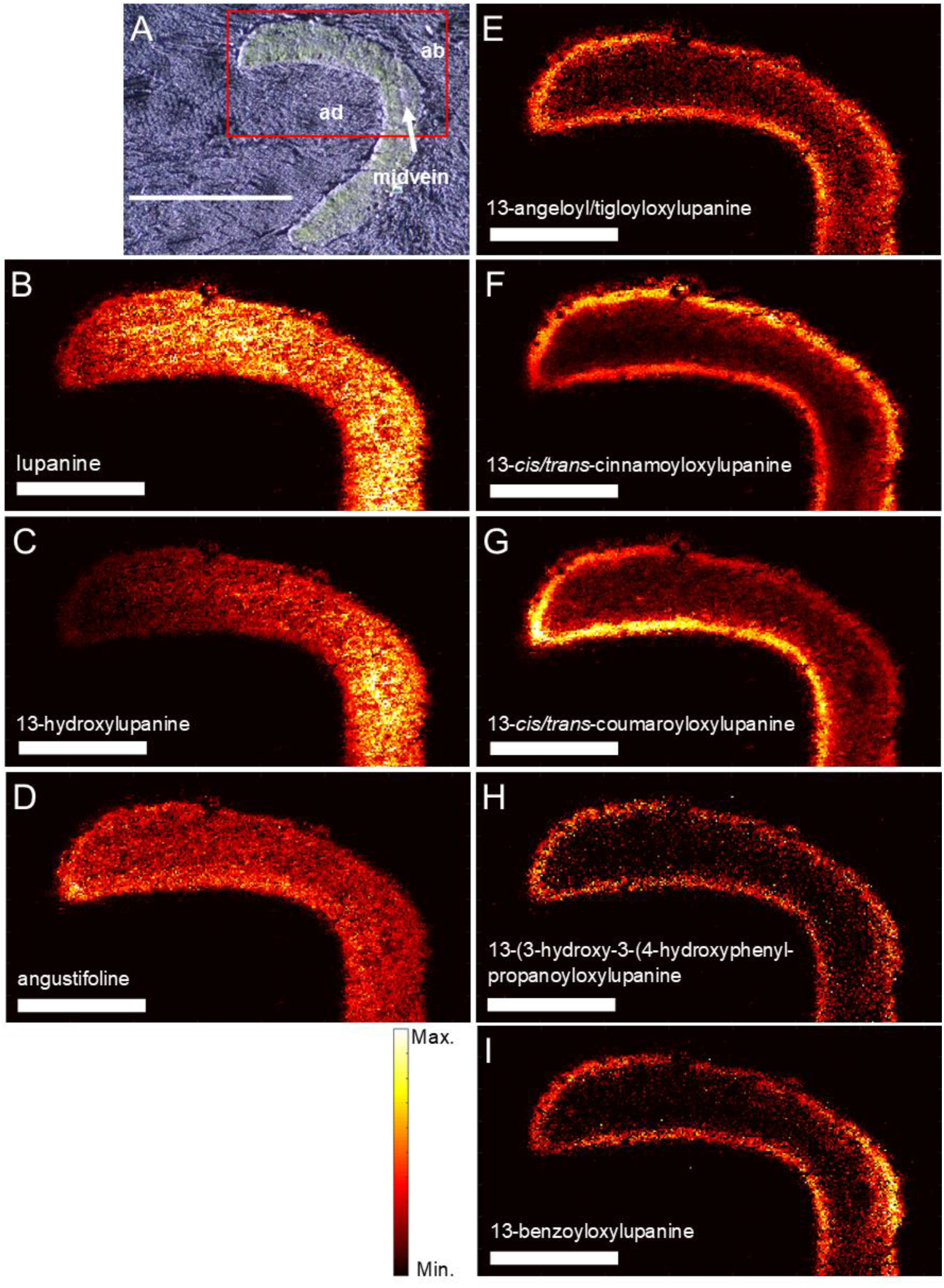
Transverse distribution of QAs in the NLL leaf. A: Bright-field microscopy image, with the red box denoting the area that was further analyzed by high-res MALDI-MSI (bar = 1 mm). ad, leaf adaxial (upper) surface; ab, leaf abaxial (lower) surface; midvein, central midvein. B–I: Individual MALDI-MS images of eight QAs at a spatial resolution of 5 μm (bar = 0.4 mm). Each image was obtained by selecting the exact mass of the protonated QA (± 5 ppm, Table S1) and normalizing by the total ion current. The color scale represents signal intensity.

#### 3.1.2 Stem

As the stem is radially symmetrical, we imaged a representative section containing all cell types from the outer epidermis to the inner pith (Fig. 2A). The core QAs were generally evenly distributed throughout the outer stem tissue, up to and including the phloem, but were not detected in the xylem or inner pith (Fig. 2, B-D). The esterified QAs seemed to be localized in the epidermis (Fig. 2, E-H).

**Fig. 2:**
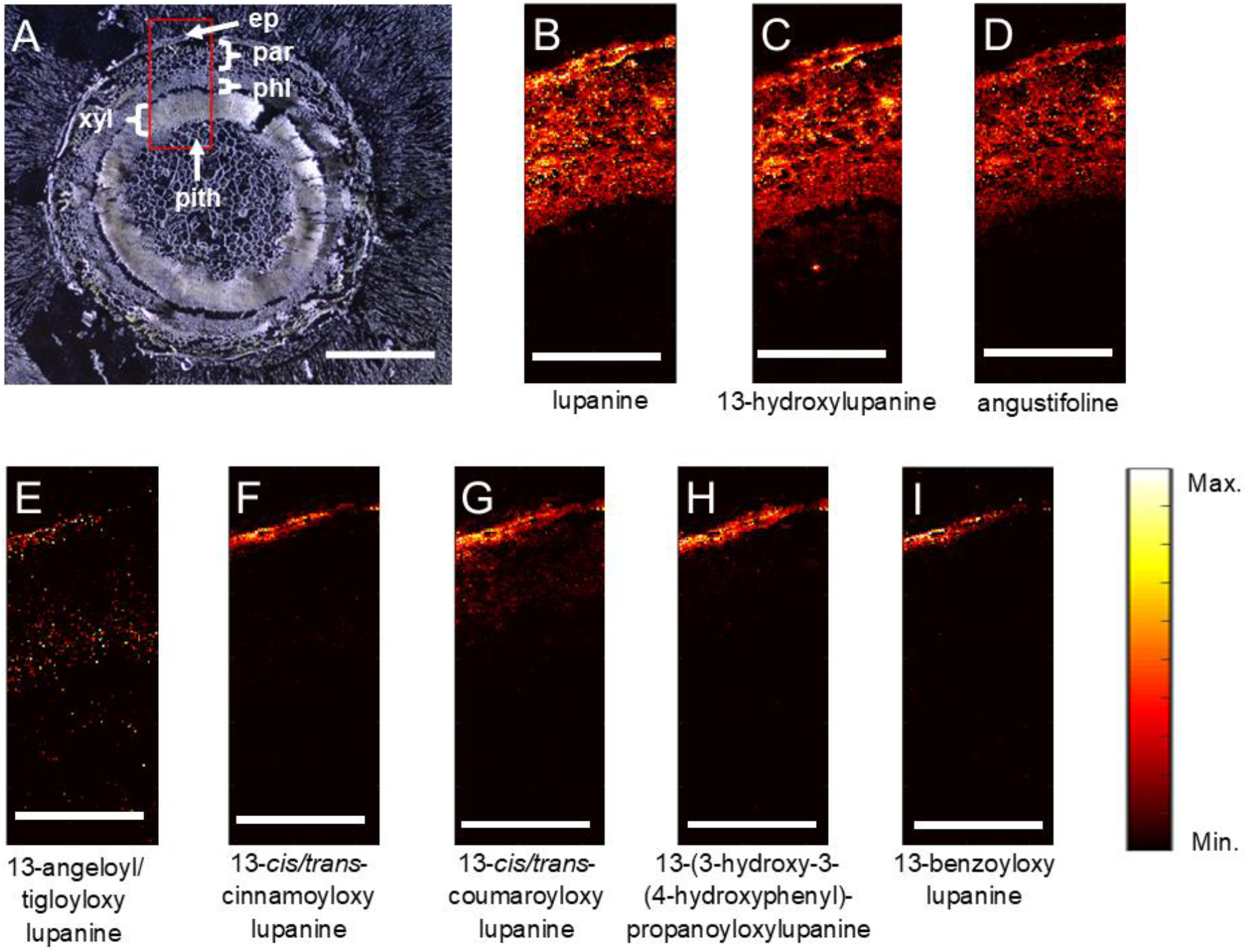
Transverse distribution of QAs in the NLL stem. A: Bright-field microscopy image, with the red box denoting the area that was further analyzed by high-res MALDI-MSI (bar = 1 mm). ep, epidermis; par, parenchyma; phl, phloem; xyl, xylem; pith, inner pith. B–I: Individual MALDI-MS images of eight QAs at spatial resolution of 5 μm (bar = 0.4 mm). Each image was obtained by selecting the exact mass of the protonated QA (± 5 ppm, Table S1) and normalizing by the total ion current. The color scale represents signal intensity.

#### 3.1.3 Pod with seed

We imaged a section of a pod with seeds inside at 30 days after anthesis (DAA), which is approximately halfway through seed development. At this point, the pods still contain relatively high levels of QAs, and these are expected to fall rapidly later in seed development (Otterbach et al., 2019). As the pod is bilaterally symmetrical, we imaged a representative portion containing the ventral suture (where the funiculus connects the pod to the seed) (Fig. 3, A and B). The core QAs displayed an even distribution throughout the pod tissue (Fig. 3, C-E), and again, the esterified QAs were predominantly in the epidermis (Fig. 3, F-J). Interestingly, we observed a high amount of core QAs as well as 13-(3-hydroxy-3-(4-hydroxyphenyl)-propanoyloxylupanine in the seed coat of the seed compared to the embryo (Fig 3, C-E and I). This could be indicative of an intermediate stage of long-distance transport, in which QAs accumulate in the seed coat before being transferred to the embryo at near maturity.

**Fig. 3:**
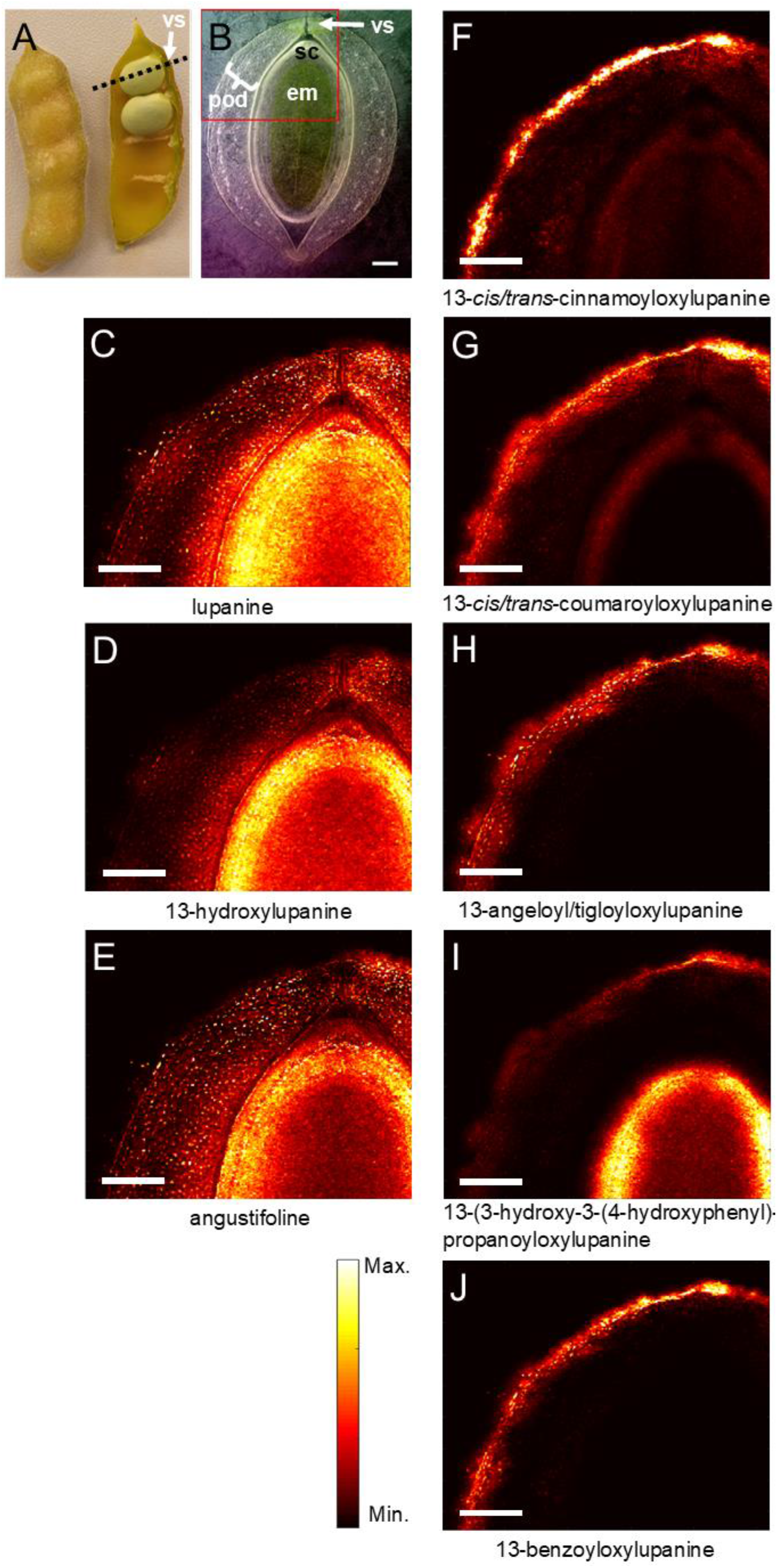
Transverse distribution of QAs in the developing NLL pod, including an enclosed seed. The pod was harvested at ~30 DAA. A: NLL pod with dotted line illustrating the analyzed transverse plane. B: Bright-field microscopy image with red box denoting the area that was further analyzed by high-res MALDI-MSI (bar = 1 mm). vs, ventral suture; pod, pod wall; sc, seed coat; em, seed embryo. C–J: Individual MALDI-MS images of QAs at 20 μm spatial resolution (bar = 1 mm). Each image was obtained by selecting the exact mass of the protonated QA (± 5 ppm, Table S1) and normalizing by the total ion current. The color scale represents signal intensity.

### 3.2. Tissue-specific localization of biosynthetic gene expression

We used laser-capture microdissection (LCM) to generate a tissue-specific RNAseq dataset from NLL leaves, stems, and developing pods. For each organ, we collected cells from as many tissues as possible, the limiting factors being the size of cells/tissues and their interconnectivity. This resulted in a different number of tissues collected for each organ, always including the epidermis (site of QA ester accumulation), the chloroplastic parenchyma (hypothesized site of QA biosynthesis), and the vasculature (involved in the long-distance transport of QAs). After RNA extraction and Illumina sequencing, we quantified the expression of transcripts using the LCM-RNAseq data and coding sequences of a previously published NLL transcriptome (Yang et al., 2017), giving a final LCM-RNAseq gene expression matrix. We have made this gene expression data available as an eFP browser (bar.utoronto.ca/efp_lupin) (Waese et al., 2017), where the expression of any NLL coding sequence can be easily visualized in transverse sections of leaves, stems, and developing pods. In addition, we have integrated the previously published RNAseq dataset (Yang et al., 2017) into the eFP browser. This previous dataset comes from eight whole NLL organs: leaf, stem, pedicel, flower, root, small pod including seeds, big pod, and big seed (Yang et al., 2017).

#### 3.2.1 Leaf

Due to the small size of the cells/tissues in NLL leaves, it was only possible to separately collect three tissues: epidermis (abaxial and adaxial together), mesophyll, and vascular bundles. The latter included midvein and transverse vascular bundles, both of which encompassed phloem, xylem, and bundle sheath cells. The QA biosynthetic genes *LDC* and *CAO* were highly expressed in the leaf epidermis (Fig. 4 A and D), and differential expression analysis revealed that *LDC* was significantly more highly expressed in epidermis compared to vasculature (log2 fold change = 6.18; *Padj* = 0.006), and that *CAO* was significantly more highly expressed in epidermis vs. mesophyll (log2 fold change = 10.44; *Padj* <.001). The expression of *LDC* and *CAO* did not differ significantly between mesophyll and vasculature. As it is possible to peel abaxial (but not adaxial) epidermis from NLL leaves, we were able to isolate abaxial epidermis RNA and verify the high expression of these QA biosynthetic genes in the leaf epidermis by qPCR. Indeed, *LDC* and *CAO* were significantly more highly expressed in the abaxial epidermis compared to the rest of the leaves (without abaxial epidermis), being 120 and 240 times more highly expressed, respectively (Fig. S1).

**Fig. 4:**
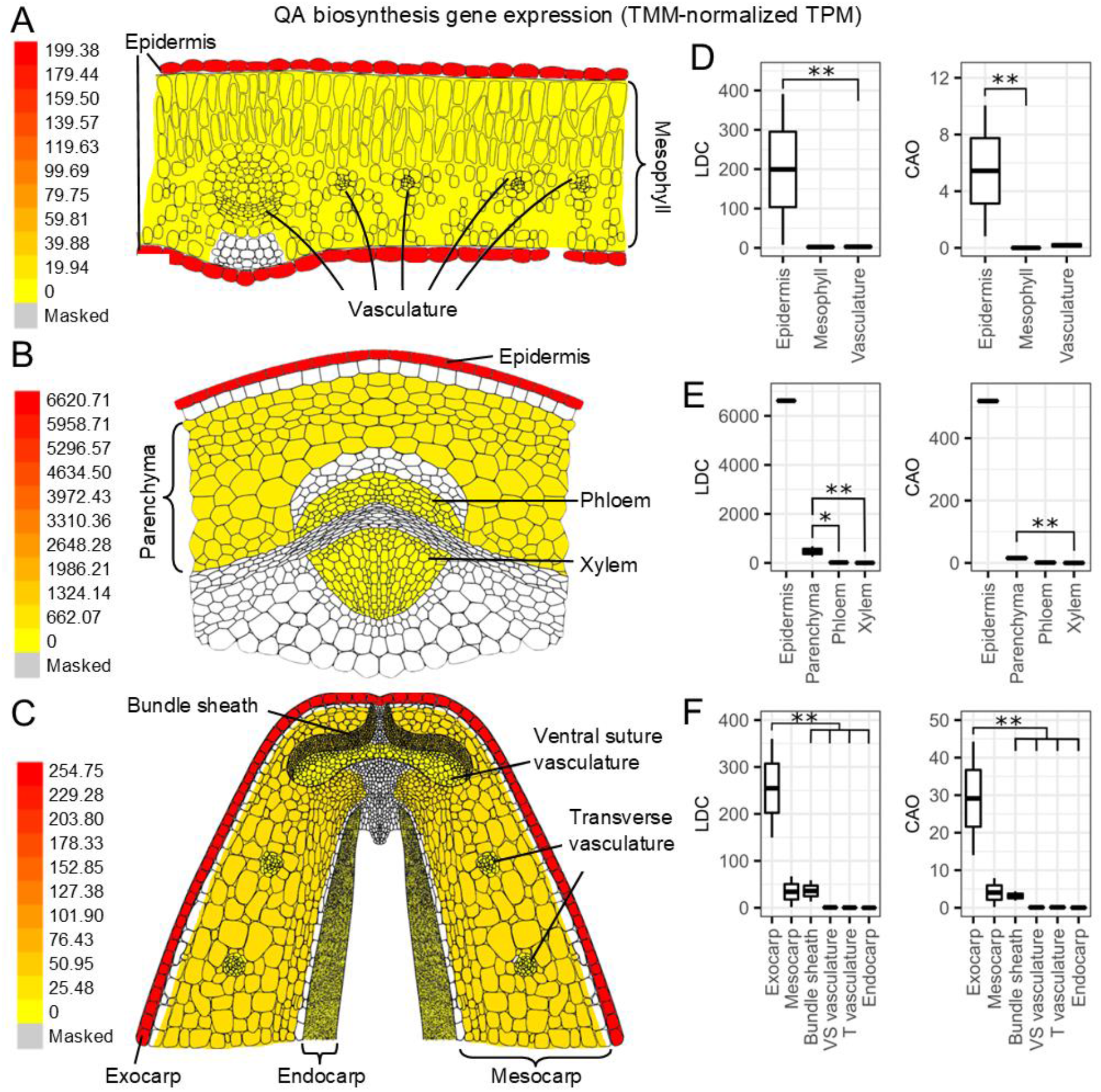
Expression of QA biosynthetic genes *LDC* and *CAO* as determined by LCM-RNAseq. A: Heat map of average *LDC* expression in leaf cells/tissues as shown on the Lupin eFP Browser, and in B: stem, and in C: developing pod. Heat map of *CAO* expression is similar and not depicted here. D: Graphical representation of *LDC* and *CAO* expression in leaf cells/tissues, and in E: stem, and in F: pod. Asterisks indicate a significant difference as determined by differential gene expression analysis: single asterisk; *Padj* < 0.05, double asterisk; *Padj* < 0.01.

#### 3.2.2 Stem

From stem tissue sections we collected epidermis, parenchyma, phloem, and xylem. While only the outer 1–3 cell layers of stem parenchyma have chloroplasts (Fig. S2), these cells were difficult to collect separately, and thus all the stem parenchyma was collected together. Additionally, it was difficult to determine the interface between phloem and vascular cambium cells, and therefore our phloem samples likely contained a small proportion of vascular cambium. Notably, however, we were able to separate the phloem from the xylem for this organ.

Unfortunately, as one RNA sample for stem epidermis failed library preparation, it was not possible to carry out statistical comparisons involving the stem epidermis. Nevertheless, *LDC* and *CAO* did have very high expression in stem epidermis compared to parenchyma, phloem, and xylem (Fig 4. B and E). Additionally, *LDC* was significantly more highly expressed in parenchyma compared to phloem and xylem (phloem: log2 fold change = 4.82, *Padj* = 0.015; xylem: log2 fold change = 6.99, *Padj* < 0.001), and *CAO* was significantly more highly expressed in parenchyma compared to xylem (log2 fold change = 9.25, *Padj* < 0.001).

#### 3.2.3 Developing pod

Due to the large size and complexity of NLL pod cells/tissue, it was possible to collect a wider variety of tissues from the developing pod. We collected exocarp (outer epidermis), mesocarp (parenchyma without vasculature), bundle sheath, vasculature at the ventral suture, vasculature transversing the mesocarp, and endocarp (inner pod layer comprised of epidermis, thin-walled parenchyma and sclerenchyma). *LDC* and *CAO* displayed high expression in the exocarp (outer epidermis) (Fig. 4 C and F), with both genes being significantly more highly expressed in this tissue compared to all others (log2 fold change = 2.86–12.66, *Padj* < 0.001) with the exception of the mesocarp.

### 3.3. Visualization of chloroplasts in the epidermis of leaves, stems, and pods

Due to the unexpected result of *LDC* and *CAO* being highly expressed in the epidermis of leaves, stems, and pods, we probed these tissues for the presence of chloroplasts, where LDC has been found to localize intracellularly (Wink and Hartmann, 1982; Bunsupa et al., 2012). To do so we imaged chlorophyll autofluorescence in the respective epidermal cells using confocal laser scanning microscopy. Indeed, the epidermis of all three organs contained chloroplasts (Fig. 5), although they were smaller and fewer in number than in the underlying cell layers (z-stacks; Supplemental Files 1–4).

**Fig. 5:**
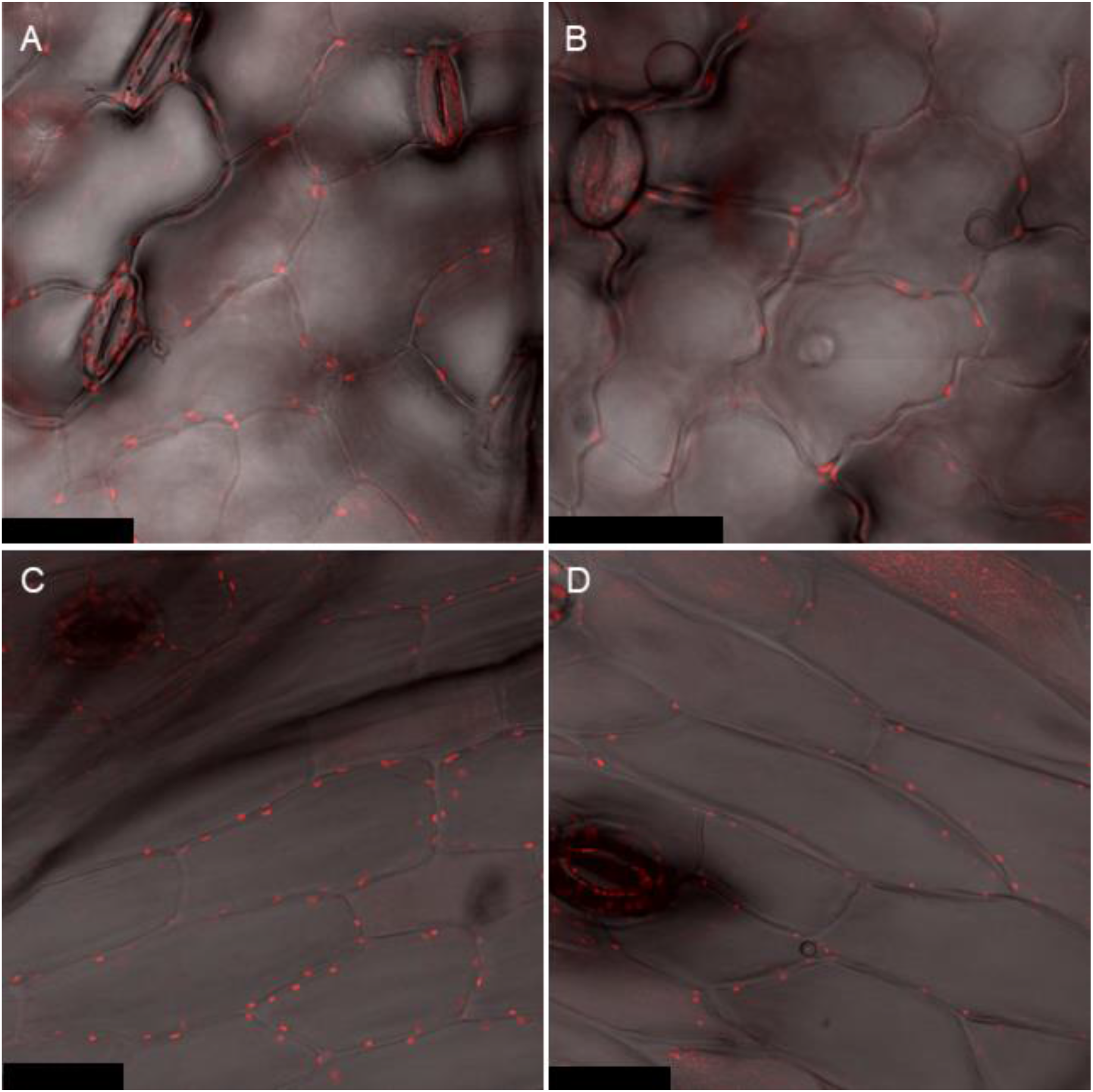
Visualization of chloroplasts in NLL epidermal cells. In each panel, chlorophyll autofluorescence (red) is overlaid on the respective bright-field microscopy image (grey). A: Leaf abaxial epidermis. B: Leaf adaxial epidermis. C: Stem epidermis. D: Developing pod epidermis (bar = 40 μm).

### 3.4 Probing the sites of QA biosynthesis in leaves via precursor feeding studies

Finally, to probe the sites of QA biosynthesis in leaves, we fed isotopically labelled L-lysine (QA precursor) and analyzed labelled QAs via tissue dissection coupled to LC-MS or via MALDI-MSI. We had initially attempted to feed the labelled L-lysine to isolated leaf epidermis, similarly to the method employed by Wink et al. (1987), who fed radiolabeled cadaverine to isolated petiole epidermis. However, we found that even after a short incubation in media (6 h), the epidermis was nearly devoid of QAs, where it had previously had high levels of QAs before the incubation (Fig. S3). As an alternative approach, we fed the isotopically labelled L-lysine to whole, detached leaves via the petiole and dissected the leaves for QA analysis at various time points after continuous feeding. We provide an illustration of how the isotopic label is incorporated into lupanine (Fig. S4); the tetracyclic QA backbone is derived from three units of L-lysine, described in detail in Mancinotti et al., (2022). As previously mentioned, it is straightforward to strip the abaxial epidermis from NLL leaves, but not the adaxial epidermis. Therefore, we collected abaxial epidermis, the remaining leaf tissue (without abaxial epidermis), and a sample of “mesophyll” (tissue from leaves without epidermis, likely containing some vasculature). To collect this last type of tissue, leaves stripped of the abaxial epidermis were frozen, and the mesophyll was isolated by carefully scraping off tissue with a scalpel blade, avoiding the adaxial epidermis.

In these experiments, the fed, labelled L-lysine was detected in all three tissue fractions, with the abaxial epidermis accumulating less than the remaining leaf tissue after 10, 18, and 24 h of feeding, and less than the mesophyll after 10 h of feeding (Fig. 6A). By contrast, the abaxial epidermis accumulated more labelled lupanine than the remaining leaf tissue or the mesophyll, significantly so after 6 and 18 h of feeding (Fig. 6B). In addition, the percentage of total lupanine that was labelled was significantly higher in the abaxial epidermis compared the remaining leaf tissue at all time points, and was also significantly higher compared to the mesophyll after 10 h of feeding (Fig. S5). Notably, after 24 h of feeding, nearly 50% of the lupanine in the abaxial epidermis was labelled (Fig. S5). Additionally, we could detect the presence of a likely side-product of the early QA pathway, ammodendrine (Mancinotti et al., 2022), in its labelled form (Fig. 6C). After 6 and 18 h of feeding, the levels of labelled ammodendrine were significantly higher in the abaxial epidermis than in the remaining leaf tissues, and at 18 h after feeding, the levels were significantly higher in the abaxial epidermis than the mesophyll (Fig. 6C). We could also detect two other labelled core QAs (13-hydroxylupanine and angustifoline), and a labelled QA ester (13-*trans*-cinnamoyloxylupanine) after 18 and 24 h of feeding (Fig. S6). However, levels were very low (barely detectable after 10 h of feeding) and for the most part did not differ significantly between the three tissue fractions (Fig. S6).

**Fig. 6:**
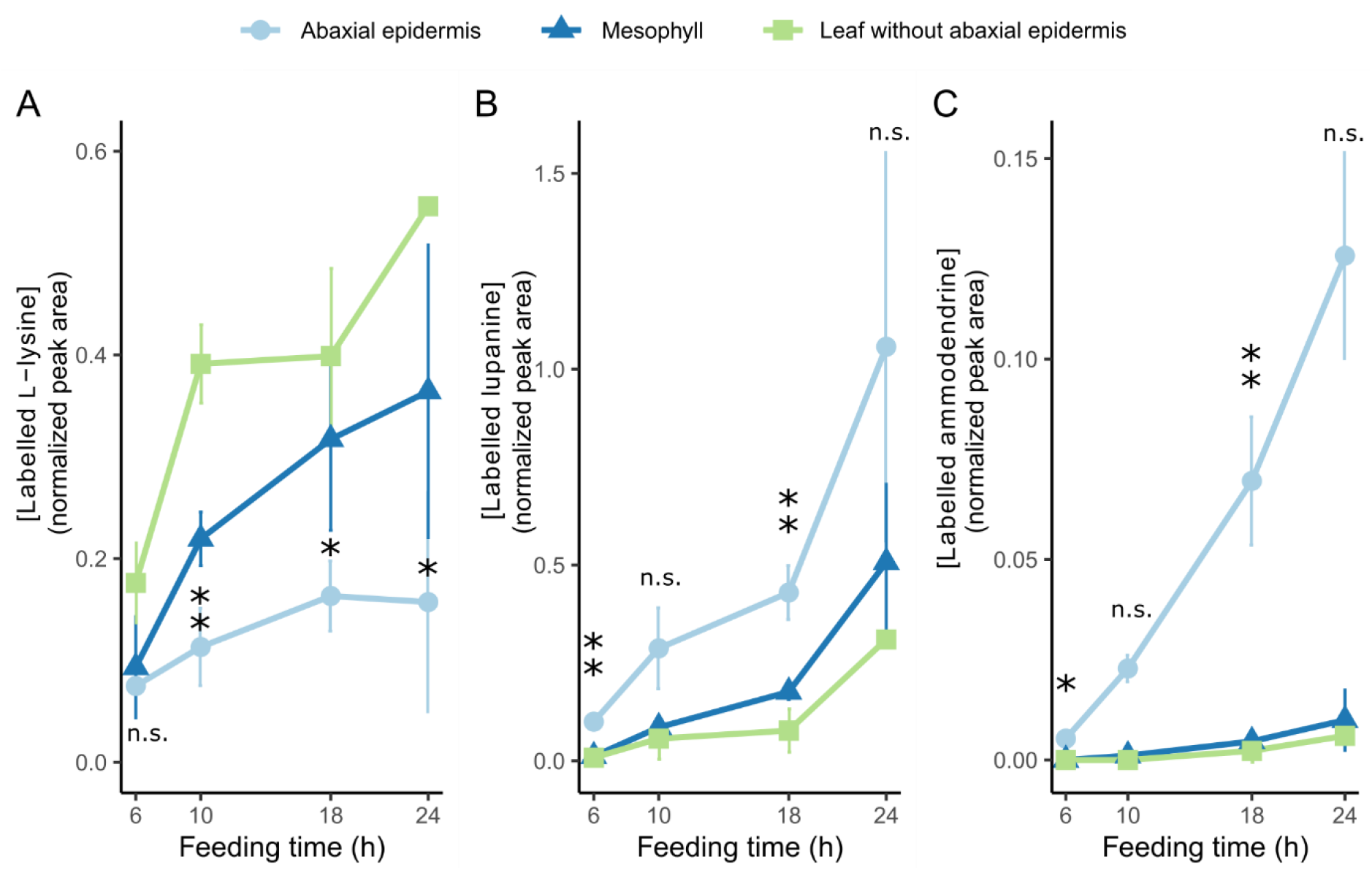
Accumulation of isotopically labelled compounds in tissue fractions of NLL leaves upon feeding with labelled L-lysine. Whole leaves were fed via the cut petiole, and leaves were dissected at four time points after the start of the feeding period (6, 10, 18, and 24 h). The respective tissue fractions (abaxial epidermis, mesophyll, and leaf without abaxial epidermis) were analyzed by LC-MS. A: Accumulation of the fed, labelled L-lysine in the tissue fractions across time points. Single asterisks represent a significant difference between abaxial epidermis and the leaf without abaxial epidermis, and double asterisks represent significant differences between abaxial epidermis and both the leaf without abaxial epidermis and the mesophyll (*Padj* < 0.05 on Tukey’s test). n.s. = not significant. B: Accumulation of the QA lupanine in the tissue fractions across time points. Significant differences are represented as described for panel A. C: Accumulation of ammodendrine (likely side product of the early QA pathway) in the tissue fractions across time points. Significant differences are represented as described for panel A, with the exception of the 6 h time point, for which a non-parametric statistical test was used (*Padj* < 0.05 on Dunn’s test).

To generate further insights on the precise sites of biosynthesis, we generated MALDI-MS images of transverse sections of the fed leaves for the early time points (6 and 10 h). While unlabeled lupanine and labelled L-lysine were distributed throughout the leaf section at 6 h and 10 h after feeding, the isotopically labelled lupanine showed a distinctive localization on the abaxial side of the leaf (Fig. 7 and Fig. S7). The accumulation began on the abaxial side of the leaf around the midrib after 6 h of feeding, and extended laterally along the abaxial side after 10 h of feeding. Negative controls were also imaged (leaves fed with unlabeled L-lysine), and they gave no signal for both the labelled L-lysine and the labelled lupanine, as expected (data not shown).

**Fig. 7:**
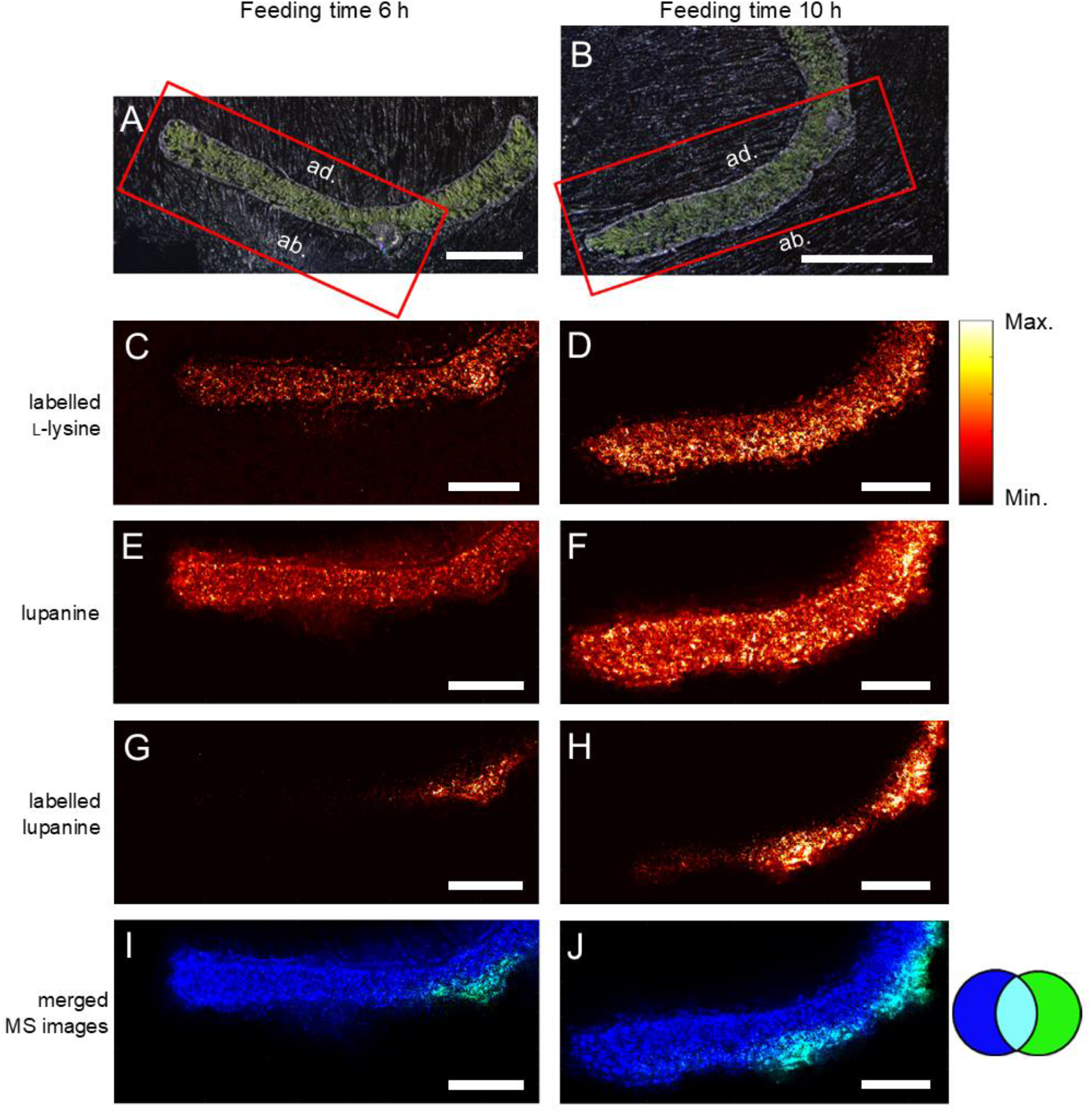
Distribution of isotopically labelled lupanine in transverse sections of NLL leaves at 6 h (left column) and 10 h (right column) after continuous feeding with isotopically labelled L-lysine. Tissue sections were analyzed by high-res MALDI-MSI at 5 μm spatial resolution. Compounds were visualized by selecting the respective *m/z* ratios of their protonated forms (± 5 ppm, Table S1) and normalizing by the total ion current. A–B: Bright-field microscopy images with red boxes representing the areas that were further analyzed (bar = 1 mm). C–D: MALDI-MSI visualization of labelled L-lysine. The color scale (right) represents signal intensity. E–F: MALDI-MSI visualization of unlabelled lupanine (same color scale). G–H: MALDI-MSI visualization of labelled lupanine (same color scale). I–J: Merged and recolored MALDI-MS images of unlabeled lupanine (blue) and labelled lupanine (green). Bar = 0.4 mm for C–J.

## 4. Discussion

Here we show that QAs in NLL are made primarily in the epidermis of QA-synthesizing organs (leaves, stems, and developing pods). Supporting evidence includes the higher expression of biosynthetic genes in the epidermis compared to other tissues (Figs. 4 and S1) as well as the larger incorporation of label from a QA precursor in the leaf epidermis compared to the rest of the leaf (Figs. 6, 7, and S5). For both lines of evidence, we collected results from two different experimental approaches. In the case of gene expression, the experimental approaches comprised LCM-RNAseq (Fig. 4) as well as manual leaf tissue dissection followed by qPCR (Fig. S1). In the case of the precursor feeding studies, we analyzed the fed leaves via manual dissection coupled to LC-MS (Figs. 6 and S5) as well as via MALDI-MSI (Fig. 7).

Our findings stand in contrast to those of Wink and Mende (1987) who concluded that QAs were made in mesophyll tissue. In their study, the radiolabeled QA intermediate cadaverine was separately fed to intact petioles of *L. polyphyllus*, the stripped petiole epidermis, and the remaining tissue (“mesophyll”). Radiolabeled lupanine was detected in intact petioles and in the “mesophyll”, but not in the epidermis. However, the authors did not report the levels of unlabeled QAs in this experiment. In our experience, when we attempted to feed isotopically labelled L-lysine to stripped leaf epidermis, the high levels of unlabeled QAs originally present became almost undetectable in the course of 6 h (Fig. S3). It is possible that a similar loss of QAs occurred in the experiment by Wink and Mende, thus preventing the detection of radiolabeled QAs in the stripped epidermis (e.g. due to loss of cell viability, rapid QA metabolism, or efficient QA export). Further experiments are needed to determine whether the robust biosynthetic capacity that we observed in the NLL epidermis can be extrapolated to other *Lupinus* species including *L. polyphyllus*.

Wink and Mende (1987) argued that epidermal cells are not able to make substantial amounts of QAs because they are usually devoid of chloroplasts—where the biosynthetic enzyme LDC localizes. Notably, in our study, we were able to observe chloroplasts in the epidermis of all biosynthetic organs of NLL (Fig. 5). While epidermal chloroplasts are often overlooked in higher plants, the presence of epidermal chloroplasts is particularly well documented for *Arabidopsis thaliana* and tobacco (Dupree et al., 1991; Barton et al., 2016). Already in 1879, it had been found that 85–95% of the 102 studied dicotyledonous species had epidermal chloroplasts, in particular, in the abaxial leaf epidermis (Stöhr, 1879). Although the function of epidermal chloroplasts is still not well understood, their size and number is smaller than those of mesophyll cells. In addition, their low chlorophyll content has led to the assumption that their photosynthetic contribution is comparatively low (Barton et al., 2016). If the activity of LDC does indeed need chloroplast localization, the required chloroplasts do exist in the epidermis of NLL biosynthetic organs. It might be of interest, however, to investigate if LDC also localizes elsewhere in epidermal cells, in particular, in other plastid types. Interestingly, functional LDC can be expressed in hairy roots of *Nicotiana tabacum*, where it is likely that it localized to the root’s leucoplasts (Bunsupa et al., 2014). The high biosynthetic capacity of the NLL epidermis may arise from localization in different types of NLL epidermal plastids or even elsewhere in epidermal cells. In any case, it is worth noting that even though LDC was initially found to localize in chloroplasts purified from whole leaf fractions of *L. polyphyllus*, the activity of LDC was 2–3 orders of magnitude lower than that of the final enzyme of lysine biosynthesis (also localized to chloroplasts) (Wink and Hartmann, 1982). This is consistent with only a fraction of the purified chloroplasts—likely the epidermal fraction—being biosynthetic with respect to QAs.

Interestingly, our precursor feeding studies coupled to MALDI-MSI suggest that QAs in NLL are synthesized in the abaxial, but not the adaxial leaf epidermis (Fig 7). This is supported by the qPCR results from manually dissected leaves, which show that very little biosynthetic gene expression remains in the leaf after peeling the adaxial epidermis off (Fig. S1). It should be noted that, in our LCM-RNAseq experiment, we collected the abaxial and the adaxial leaf epidermis together. Thus, the depiction of *LDC* expression in both epidermal layers in Fig. 4A is due to our inability to discriminate between these layers in this experiment (rather than a representation of our running hypothesis). It is not clear why QA biosynthesis should take place mostly or only in the abaxial leaf epidermis. Leaf development is subject to strict regulation that establishes and maintains the abaxial–adaxial axis (Manuela and Xu, 2020).

Since very little is known about the regulation of QA biosynthesis, further studies are needed to uncover any connections between abaxial-adaxial axis establishment and QA biosynthesis.

Our new finding that QAs are made primarily in the epidermis of QA-synthesizing organs has important implications for the elucidation for the entire QA pathway, for which only the first two genes are known (Mancinotti et al., 2022). The LCM-RNAseq dataset generated in this study is a valuable resource in the discovery of QA biosynthesis genes, which are likely also highly expressed in the epidermis, at least until the formation of lupanine. Indeed, in our feeding experiments, we observed much higher signals for labelled lupanine than for any other labelled QA (Figs. 6 and S6). It should be noted that our feeding experiments lasted at most 24 hours, and thus, it is possible that longer feeding experiments would lead to a higher conversion towards 13-hydroxylupanine and its esters. Notably, MALDI-MSI revealed that QA esters localize primarily to the epidermis of QA-synthesizing organs (Figs. 1, 2, and 3), suggesting (but not yet confirming) that QA esters are synthesized in the epidermis of these organs and remain there. By contrast, the core QAs seem to be transported to the underlying cell layers, as observed for labelled lupanine in the MALDI-MS images of precursor-fed leaves (Fig. 7). Eventually, the core QAs must be transported into the phloem (Lee et al., 2006) for their translocation to the seeds (Otterbach et al., 2019). We present our new working hypothesis in Fig. 8. The molecular mechanisms by which this transport pathway takes place remain to be discovered. A full understanding of QA biosynthesis and transport in NLL may allow for the targeted removal of QAs from the seeds to increase the nutritional value of the high-protein seeds.

**Fig. 8:**
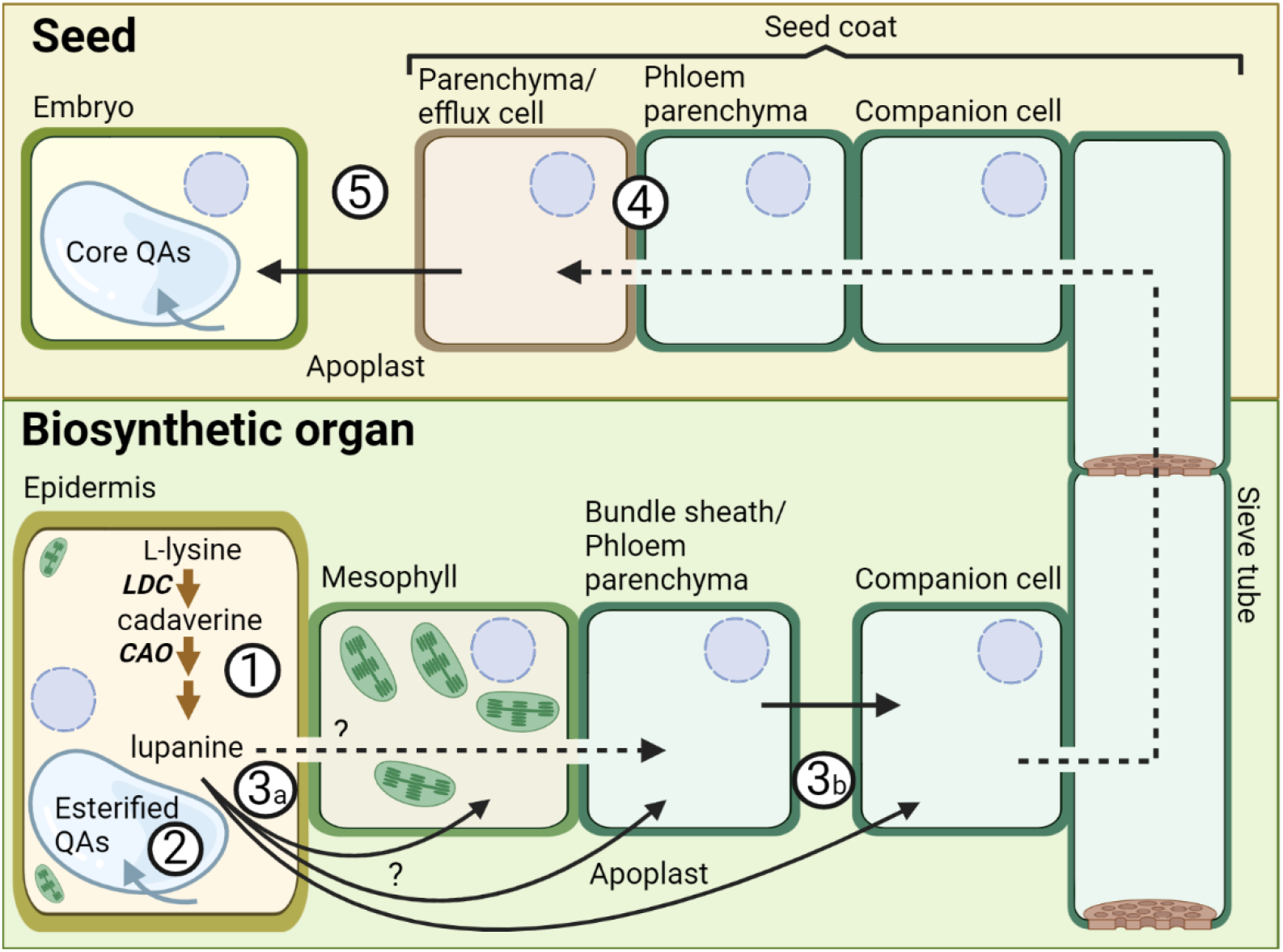
Spatial model for QA biosynthesis and transport. The aerial epidermis is a major site of QA biosynthesis, at least until the formation of lupanine (1) (pathway sub-cellular localization is yet to be defined). The esterified QAs accumulate in the epidermis, likely within the vacuole (2) (Mende and Wink, 1987). The core QAs (e.g. lupanine) must move from the epidermis to the phloem-associated cells for long-distance transport to the seeds. It is yet to be shown whether this is via a symplastic or apoplastic route, or both (3a). The apoplastic phloem loading mechanism depicted (3b) is generally found in Fabaceae species (although not yet confirmed for NLL) (Bourquin et al., 1990; Wimmers and Turgeon, 1991). After long-distance transport through the phloem, the QAs likely enter the seed coat cells through symplastic connections with the vasculature (4) (van Dongen et al., 2003; Tegeder, 2014). Finally, an apoplastic transport step is required for the QAs to accumulate in embryo cells (5) (Tegeder, 2014). Dotted black arrows represent symplastic transport, solid black arrows represent apoplastic transport, blue arrows represent transport into the vacuole, and brown arrows represent QA biosynthesis steps. Created with BioRender.com.

While the core QAs accumulate evenly across tissues and are transported long-distance to the seeds, the esterified QAs are found primarily in the epidermis of biosynthetic organs (Figs 1, 2 and 3). This suggests that the esterified QAs play a distinct biological role in the above-ground epidermis. QAs have long been thought to play a role in plant protection, particularly against insect pests. Interestingly, the QA ester 13-angeloyl/tigloyloxylupanine has been reported to be a more effective antimicrobial agent than the core QA lupanine (Wink, 1984). In addition, both 13-angeloyl/tigloyloxylupanine and 13-*cis/trans*-coumaroyloxylupanine were found to be highly deterrent against spruce budworm, whereas the core QAs were not (Bentley et al., 1984). The study of the molecular mechanisms for biosynthesis and storage of QA esters in the NLL epidermis will help elucidate their distinct physiological roles.

### Conclusion

Here we show for the first time that the QAs in NLL are mainly synthesized in the epidermis of biosynthetic organs (leaves, stems, and developing pods). While the core QAs are transported to other tissues and ultimately translocated to the seeds, the esterified QAs accumulate to high levels in the epidermis of biosynthetic organs. Furthermore, we provide a valuable tissue-specific LCM-RNAseq dataset, which we have made available as a Lupin eFP Browser together with a previously generated organ-specific RNAseq dataset. Our work uncovers new aspects of QA biosynthesis and transport, setting the stage for the discovery of the underlying molecular mechanisms.

## 5. Acknowledgements

We would like to thank the Center for Advanced Bioimaging (CAB) Denmark, and particularly Sebastian Kjeldgaard-Nintemann, for providing instrumentation and guidance. We also thank Javiera Aravena-Calvo (drawmyscience.com) for providing scientific illustrations, Oliver Gericke for advice regarding RNAseq data analysis, and Nikola Micic for assistance in matrix sublimation experiments. The project was funded by the European Union’s Horizon 2020 research and innovation programme (Marie Sklodowska-Curie grant agreement 846089) and by the Novo Nordisk Foundation (NovoCrops, project NNF2019OC53580). MDBBL was funded by the European Union’s Horizon 2020 research and innovation program (MIAMi project grant agreement 814645). Support from the Carlsberg Foundation and The Danish Council for Independent Research|Medical Sciences (Grant DFF-4002-00391) for the applied MALDI-MSI instrumentation is gratefully acknowledged.

## 6. Competing interests

None declared.

## 7. Author contributions

KMF and FG-F conceived the research plan. KMF and MDBL carried out the experiments, analyzed the data, and prepared the figures. NJP, EE and AP generated the Lupin eFP Browser. AS, CJ, and HHN-E provided supervision and contributed with data interpretation. FG-F coordinated and supervised the project. KMF wrote the manuscript with input from all authors.

## 8. Data availability

The MSI data generated during this study are available on METASPACE (https://metaspace2020.eu/project/lupinus_angustifolius_Frick-et-al), and the RNAseq data generated during this study are available on the NCBI Sequence Read Archive (SRA) (BioProject PRJNA943121).

## 9. Supporting Information

**Table S1:** List of compounds examined via MALDI-MSI and/or LC-MS.

**Fig. S1:** Expression of QA biosynthetic genes in leaf abaxial epidermis compared to the rest of the leaf without abaxial epidermis, as determined by qPCR.

**Fig. S2:** Fluorescence microscopy image of a cross section of an NLL stem.

**Fig. S3:** Normalized relative peak area of quinolizidine alkaloids (QAs) in abaxial epidermis of narrow-leafed lupin leaves, after 0 h and 6 h incubation in media (B5 medium, 2% sucrose and 1-5mM lysine) as determined by LC-MS.

**Fig. S4:** Incorporation of the isotopically labelled L-lysine used in the feeding experiments into the tetracyclic QA backbone.

**Fig. S5:** Percentage of total amount of lupanine with isotopic label in tissue fractions of NLL leaves upon feeding with labelled L-lysine.

**Fig. S6:** Accumulation of isotopically labelled alkaloids in tissue fractions of NLL leaves upon feeding with labelled L-lysine.

**Fig. S7:** Distribution of isotopically labelled L-lysine and PC(34:2) NLL transverse leaf sections at 6 h and 10 h after feeding with isotopically labelled L-lysine.

**Supplemental File 1:** Chlorophyll autofluorescence (red) in NLL leaf abaxial epidermis and underlying cell layers (z-stack).

**Supplemental File 2:** Chlorophyll autofluorescence (red) in NLL leaf adaxial epidermis and underlying cell layers (z-stack).

**Supplemental File 3:** Chlorophyll autofluorescence (red) in NLL stem epidermis and underlying cell layers (z-stack).

**Supplemental File 4:** Chlorophyll autofluorescence (red) in NLL developing pod epidermis and underlying cell layers (z-stack).

## Notes

### Competing Interest Statement

The authors have declared no competing interest.

