## Supporting Information for "The aerial epidermis is a major site of quinolizidine alkaloid biosynthesis in narrow-leafed lupin"

**Table S1**: List of compounds examined via MALDI-MSI and/or LC-MS. Compound formulas, structures, and associated *m/z* values for detection are shown. *Only one of these isomer pairs accumulates in NLL, we did not determine which one. ‡These isomers could be differentiated via LC-MS as described in (Otterbach et al., 2019).

| Compound | Chemical formula (analyzed form) | Chemical structures of QAs, precursors or intermediates | *m/z* |
|---|---|---|---|
| $[PC(34:2) + K]^+$ | | | 796.5253 |
| lupanine | $C_{15}H_{25}N_2O^+$ $[M+H]^+$ | 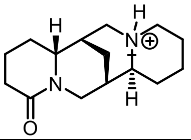 | 249.1961 |
| Isotopically labelled lupanine | $^{13}C_{15}H_{25}{}^{15}N_2O^+$ $[M+H]^+$ | As above | 266.2405 |
| 13-hydroxylupanine | $C_{15}H_{25}N_2O_2^+$ $[M+H]^+$ | 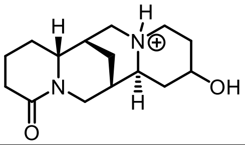 | 265.1911 |
| Isotopically labelled 13-hydroxylupanine | $^{13}C_{15}H_{25}{}^{15}N_2O_2^+$ $[M+H]^+$ | As above | 282.2354 |
| angustifoline | $C_{14}H_{23}N_2O^+$ $[M+H]^+$ | 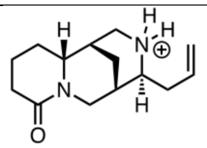 | 235.1805 |
| Isotopically labelled angustifoline | $^{13}C_{14}H_{23}{}^{15}N_2O^+$ $[M+H]^+$ | As above | 251.2215 |
| 13-angeloyl/ tigloyloxylupanine* | $C_{20}H_{31}N_2O_3^+$ $[M+H]^+$ | 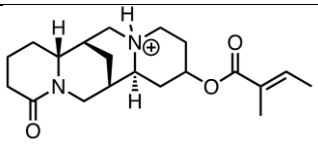 | 347.2329 |
| 13-*cis*-cinnamoyloxylupanine‡ | $C_{24}H_{31}N_2O_3^+$ $[M+H]^+$ | 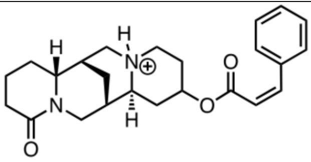 | 395.2329 |
| 13-*trans* - cinnamoyloxylupanine‡ | $C_{24}H_{31}N_2O_3^+$ $[M+H]^+$ | 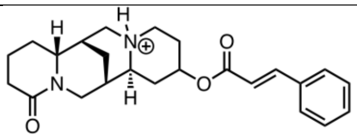 | 395.2329 |
| Isotopically labelled 13-*trans* - cinnamoyloxylupanine | $^{13}C_{15}C_9H_{31}{}^{15}N_2O_3^+$ $[M+H]^+$ | As above | 412.2773 |
| 13-*cis/trans*-coumaroyloxylupanine* | $C_{24}H_{31}N_2O_4^+$ $[M+H]^+$ | 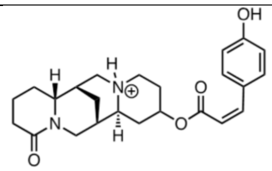 | 411.2278 |

| | | | |
|---|---|---|---|
| 13-(3-hydroxy-3-(4-hydroxyphenyl)-propanoyloxylupanine | $C_{24}H_{33}N_2O_5^+$ $[M+H]^+$ | 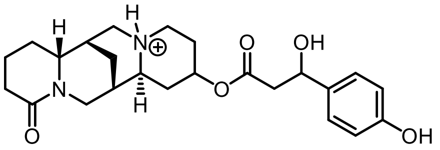 | 429.2384 |
| 13-benzoyloxylupanine | $C_{22}H_{29}N_2O_3^+$ $[M+H]^+$ | 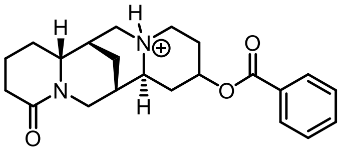 | 369.2173 |
| ammodendrine | $C_{12}H_{21}N_2O^+$ $[M+H]^+$ | 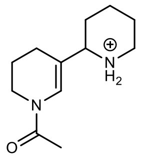 | 209.1648 |
| Isotopically labelled ammodendrine | $^{13}C_{10}C_2H_{21}{}^{15}N_2O^+$ $[M+H]^+$ | As above | 221.1925 |
| L-lysine (examined via MALSI-MSI) | $C_6H_{15}N_2O_2^+$ $[M+H]^+$ | 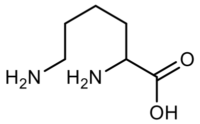 | 147.1128 |
| Isotopically labelled L-lysine (examined via MALSI-MSI) | $^{13}C_6H_{15}{}^{15}N_2O_2^+$ $[M+H]^+$ | As above | 155.1270 |
| L-lysine (examined via LC-MS) | $C_6H_{12}NO_2^+$ $[M – NH_3]^+$ | Structure undetermined | 130.0863 |
| Isotopically labelled L-lysine (examined via LC-MS) | $^{13}C_6H_{12}{}^{15}NO_2^+$ $[M – NH_3]^+$ | As above | 137.1034 |

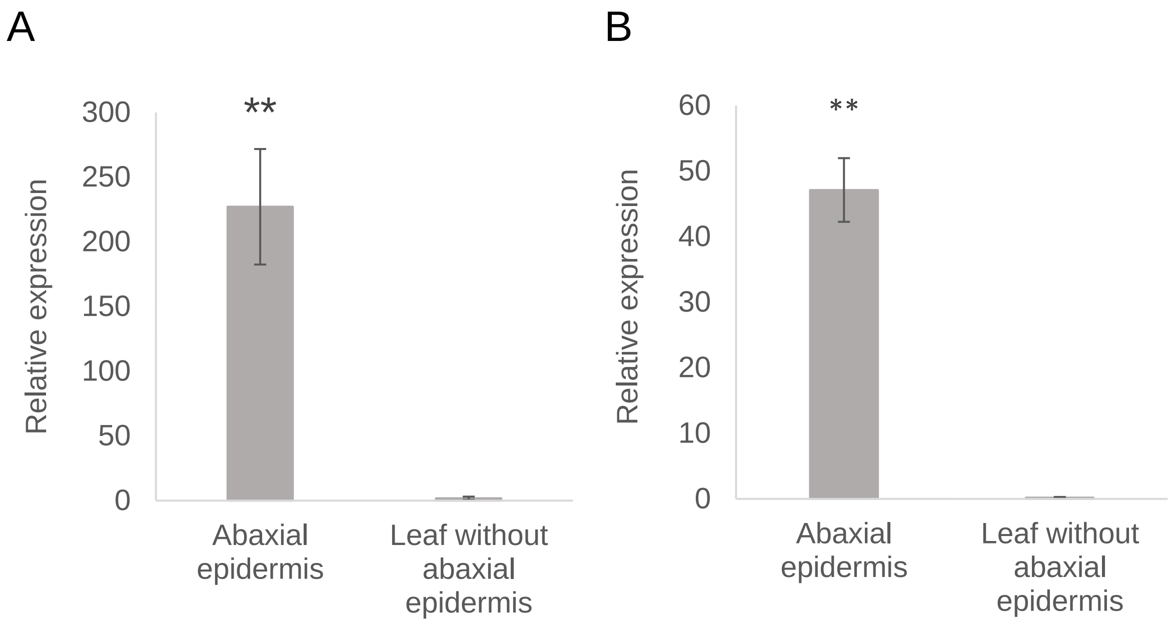


**Fig. S1:** Expression of QA biosynthetic genes in leaf abaxial epidermis compared to the rest of the leaf without abaxial epidermis, as determined by qPCR. A: *LDC* expression. B: *CAO* expression. Columns represent the mean of three biological replicates, and error bars represent the standard deviation. Asterisks indicate significant differences ($P < 0.001$), as determined by Student's t-test.

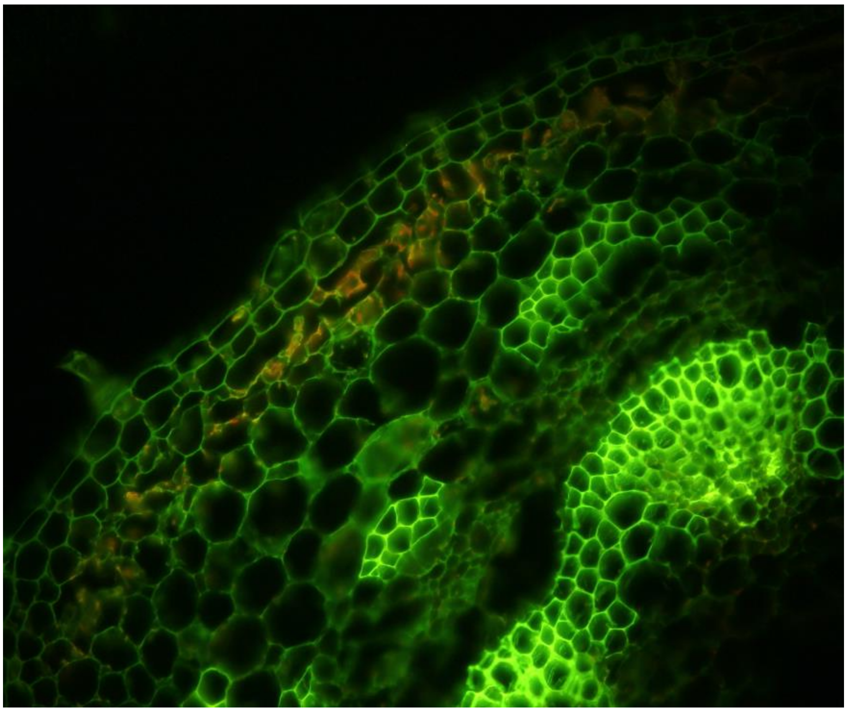

**Fig. S2:** Fluorescence microscopy image of a cross section of an NLL stem. Chlorophyll auto fluorescence (red) can be seen in the first 1-3 outer layers of the stem parenchyma. Excitation: band pass 450-490 nm; emission: long pass 515 nm.

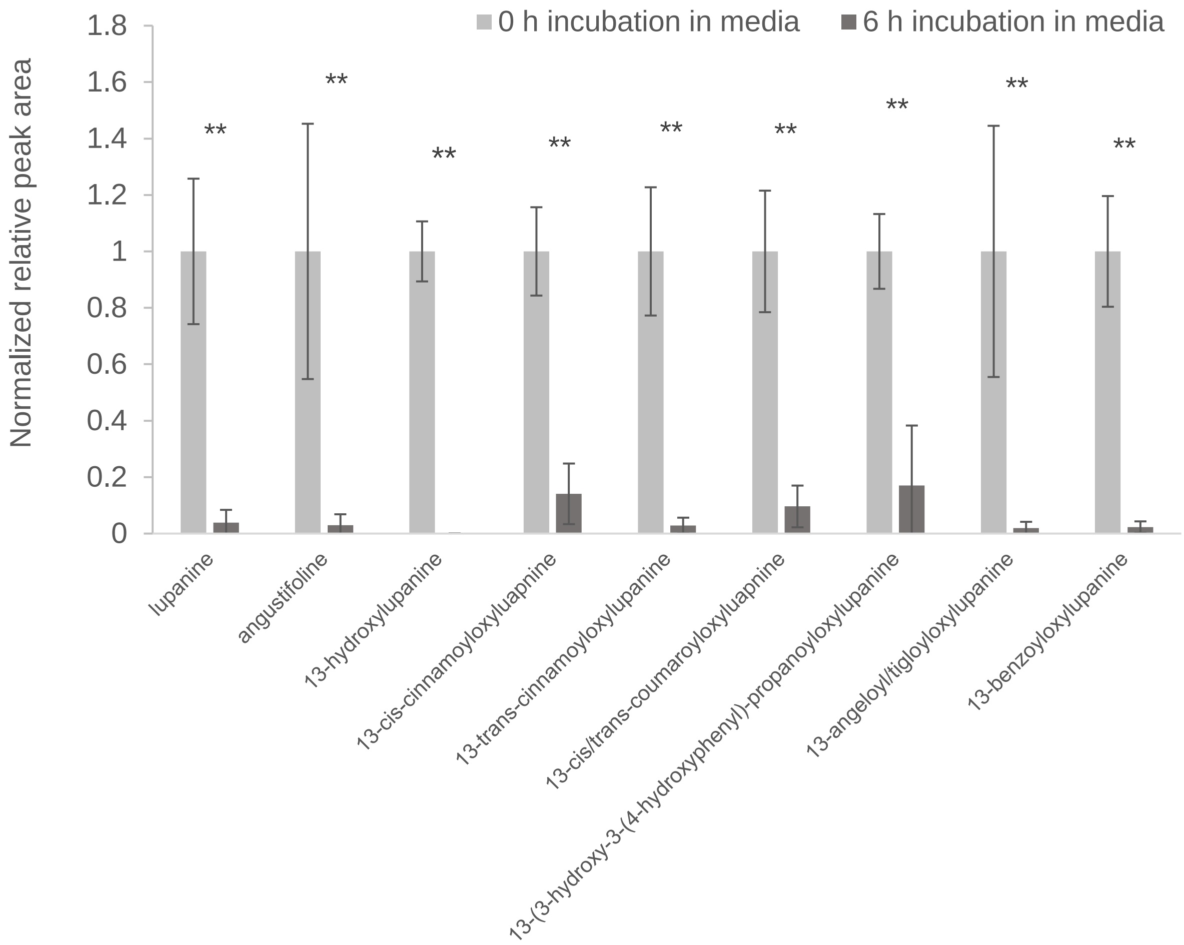


**Fig. S3:** Normalized relative peak area of quinolizidine alkaloids (QAs) in abaxial epidermis of narrow-leafed lupin leaves, after 0 h and 6 h incubation in media (B5 medium, 2% sucrose and 1–5mM lysine) as determined by LC-MS. QAs were identified by their *m/z* values (Table S1). Peak areas were determined relative to that of caffeine (internal standard) and were normalized to the corresponding values at 0 h. Asterisks indicate significant differences ($P < 0.001$) between levels of individual QAs at 0 h and 6 h as determined by Student's t-test.

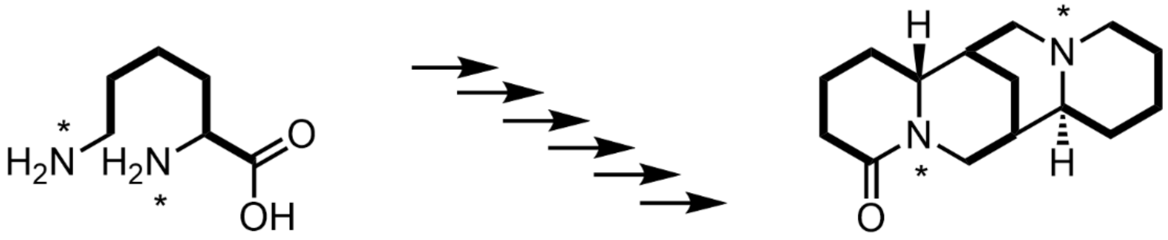


Labelled L-lysine
$^{13}C_6H_{14}{}^{15}N_2O_2$

Labelled lupanine
$^{13}C_{15}H_{24}{}^{15}N_2O$

**Fig. S4:** Incorporation of the isotopically labelled L-lysine used in the feeding experiments into the tetracyclic QA backbone (derived from three units of L-lysine). Lupanine is depicted as an example. The bonds in bold indicate the presence and incorporation of $^{13}C$, and the asterisks indicate the presence and incorporation of $^{15}N$. A compilation of previous precursor feeding experiments has been presented by Mancinotti *et al.* (2022).

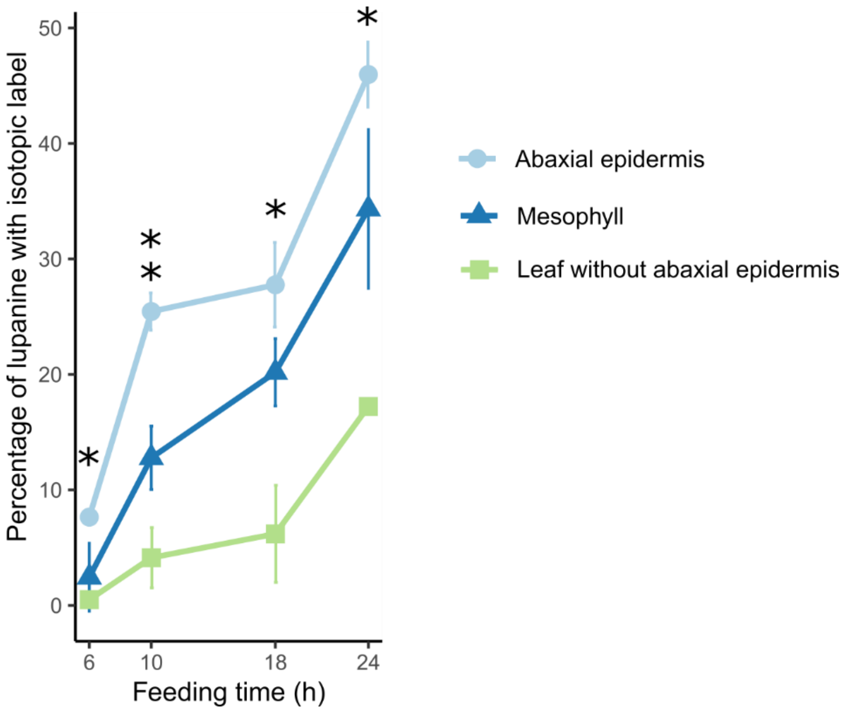


**Fig. S5**: Percentage of total amount of lupanine with isotopic label in tissue fractions of NLL leaves upon feeding with labelled L-lysine. Single asterisks represent a significant difference between abaxial epidermis and the remaining leaf tissue, and double asterisks represent significant differences between abaxial epidermis and both the remaining leaf tissue and the mesophyll (*Padj* < 0.05 on Tukey's test).

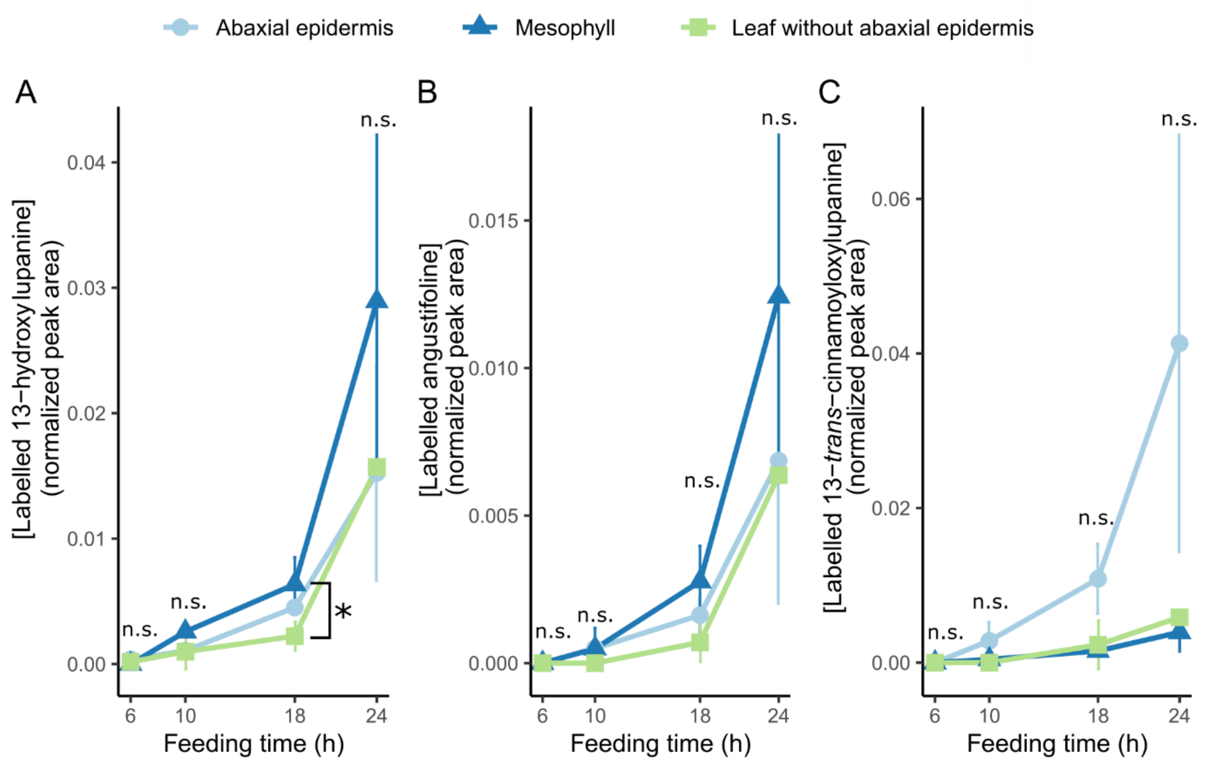


**Fig. S6**: Accumulation of isotopically labelled alkaloids in tissue fractions of NLL leaves upon feeding with labelled L-lysine. Whole leaves were fed via the cut petiole, and leaves were dissected at four time points after the start of the feeding period (6, 10, 18, and 24 h). The respective tissue fractions (abaxial epidermis, mesophyll, and leaf without abaxial epidermis) were analyzed by LC-MS. A: Accumulation of the core QA 13-hydroxylupanine in the tissue fractions across time points. B: Accumulation of the core QA angustifoline in the tissue fractions across time points. C: Accumulation of the QA ester 13-*trans*-cinnamoyloxylupanine in the tissue fractions across time points. The asterisk represents a significant difference (*Padj* < 0.05 on Tukey's test). n.s. = not significant.

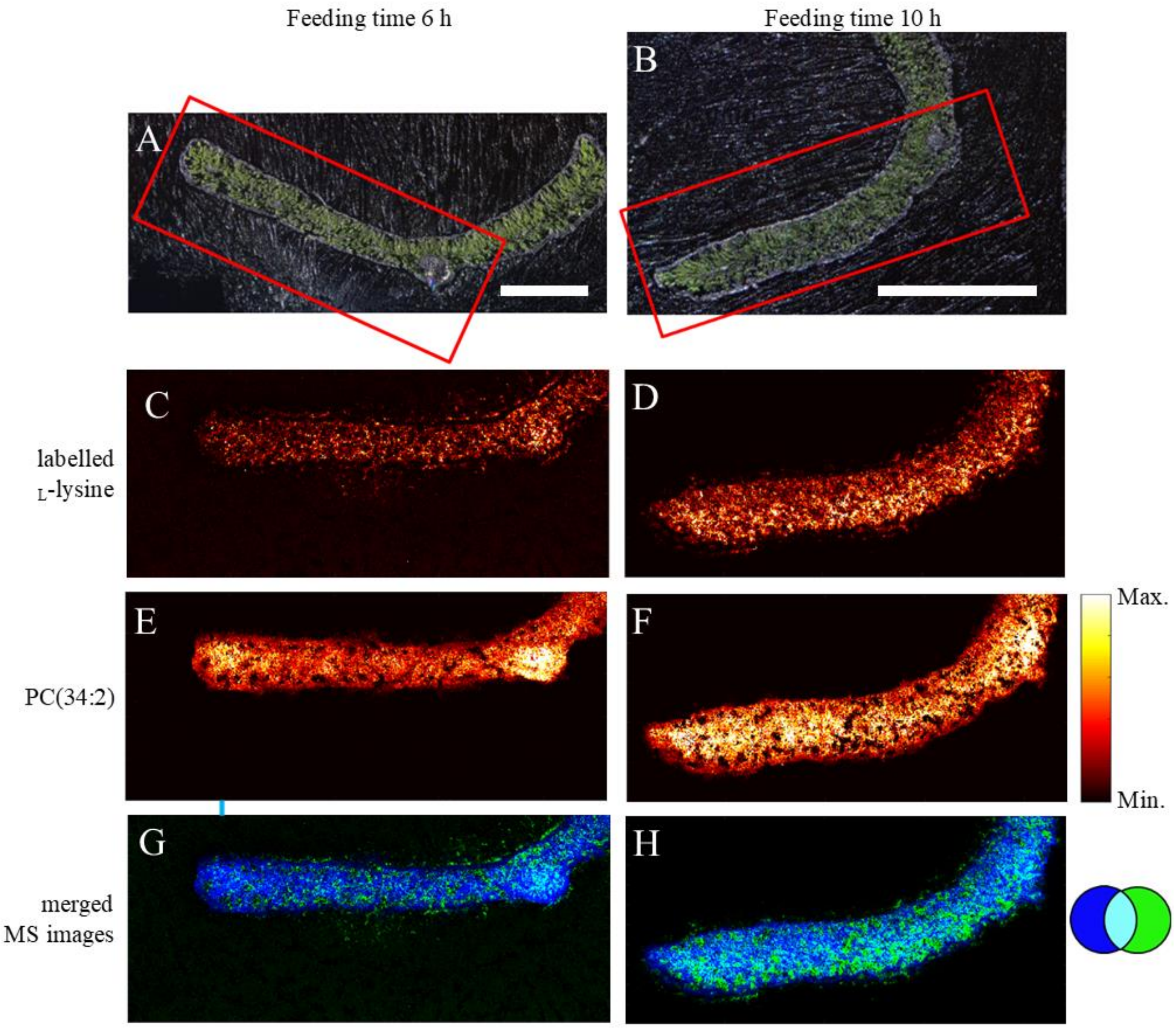


**Fig. S7:** Distribution of isotopically labelled L-lysine and PC(34:2) NLL transverse leaf sections at 6 h and 10 h after feeding with isotopically labelled L-lysine. A–B: Microscope pictures with red box denoting the area that was analyzed (Bar = 1 mm). C–F: MALDI-MS images of labelled L-lysine or PC(34:2) at 5 µm spatial resolution. G–H: Merged MALDI-MS images of labelled L-lysine (green) and PC(34:2) (blue) showing distribution of labelled L-lysine throughout the entire leaf.

### Supplemental Files:

Supplemental File 1: Chlorophyll autofluorescence (red) in NLL leaf abaxial epidermis and underlying cell layers (z-stack).

Supplemental File 2: Chlorophyll autofluorescence (red) in NLL leaf adaxial epidermis and underlying cell layers (z-stack).

Supplemental File 3: Chlorophyll autofluorescence (red) in NLL stem epidermis and underlying cell layers (z-stack).

Supplemental File 4: Chlorophyll autofluorescence (red) in NLL developing pod epidermis and underlying cell layers (z-stack).
